# 20-hydroxyecdysone (20E) signaling regulates amnioserosa morphogenesis during *Drosophila* dorsal closure: Ecdysone receptor modulates gene expression in a complex with the AP-1 component, Jun

**DOI:** 10.1101/2021.01.14.426760

**Authors:** Byoungjoo Yoo, Hae-yoon Kim, Xi Chen, Weiping Shen, Ji Sun Jang, Olga Cormier, Charles Krieger, Bruce Reed, Nicholas Harden, Simon Ji Hau Wang

**Affiliations:** Department of Molecular Biology and Biochemistry, Simon Fraser University, 8888 University Drive, Burnaby, BC, V5A 1S6, Canada; Department of Biology, University of Waterloo, 200 University Avenue West, Waterloo, ON, N2L 3G1, Canada; Department of Biomedical Physiology and Kinesiology, Simon Fraser University, 8888 University Drive, Burnaby, BC, V5A 1S6, Canada

**Keywords:** 20-hydroxyecdysone, Ecdysone receptor, Jun, *zipper*, amnioserosa, dorsal closure

## Abstract

Steroid hormones influence diverse biological processes throughout the animal life cycle, including metabolism, stress resistance, reproduction, and lifespan. In insects, the steroid hormone, 20-hydroxyecdysone (20E), is the central regulator of molting and metamorphosis, and has been shown to play roles in tissue morphogenesis. For example, amnioserosa contraction, which is a major driving force in *Drosophila* dorsal closure (DC), is defective in embryos mutant for 20E biosynthesis. Here, we show that 20E signaling modulates the transcription of several DC participants in the amnioserosa and other dorsal tissues during late embryonic development, including the *zipper* locus, which encodes for non-muscle myosin II heavy chain. Canonical 20E signaling typically involves the binding of Ecdysone receptor (EcR) and Ultraspiracle heterodimers to ecdysone-response elements (EcREs) within the promoters of ecdysone-responsive genes to drive their expression. During DC, we provide evidence that 20E signaling instead acts in parallel to the JNK cascade via a direct interaction between EcR and the AP-1 component, Jun, which together binds to genomic regions containing AP-1 binding sites but no EcREs to control gene expression. Our work demonstrates a novel mode of action for 20E signaling in *Drosophila* that likely functions beyond DC, and may provide further insights into mammalian steroid hormone receptor interactions with AP-1.

## INTRODUCTION

Dorsal closure (DC) of the *Drosophila* embryo is a developmental wound-healing event in which a hole in the dorsal epidermis, occupied by a transient epithelium, the amnioserosa, is closed by migration of the epidermal flanks (reviewed in Harden, 2002). DC serves as a paradigm for morphogenetic events where tissues are brought together and fused, including the vertebrate processes of embryonic neural tube closure and palate fusion. A recurring finding in studies of wound healing and developmental epithelial closures is that cells occupying the hole contribute to closure by contracting in response to signaling from the wound margin by TGF-β superfamily ligands (reviewed in Belacortu and Paricio, 2011). This mechanism is conserved in DC where the dorsal-most epidermal (DME) cells secrete Decapentaplegic (Dpp), a TGF-β superfamily ligand that activates a signaling pathway in the amnioserosa through the receptors Thickveins (Tkv) and Punt, which are required for correct amnioserosa morphogenesis (Fernandez et al., 2007; Wada et al., 2007; Zahedi et al., 2008). Recent studies suggest that autonomous contraction of the amnioserosa alone can drive DC and it is of interest to know how this is initiated (Pasakarnis et al., 2016; Wells et al., 2014). One way that synchronized contraction of the amnioserosa cells could be achieved is through an autocrine signaling process in which the amnioserosa cells produce a ligand that induces their own contraction. In a search for such a pathway downstream of Dpp in the amnioserosa, we considered signaling by the steroid hormone 20-hydroxyecdysone (20E). The amnioserosa is a major source of 20E during embryogenesis, and mutants of the Halloween group of genes, which encode enzymes in the 20E biosynthetic pathway, display DC defects (Chavez et al., 2000; Giesen et al., 2003; Kozlova and Thummel, 2003; Niwa et al., 2010; Ono et al., 2006).

Canonical 20E signaling involves the binding of Ecdysone receptor (EcR) and Ultraspiracle (Usp) heterodimers to ecdysone-response elements (EcREs) to promote gene expression (Dobens et al., 1991; Yao et al., 1993). We show here that 20E modulates gene expression in the amnioserosa and other dorsal tissues in a novel manner. Key DC participants in the DME cells and amnioserosa are transcribed in response to a JNK MAPK cascade operating through the AP-1 transcription factor, which consists either as a homodimer of Jun or a heterodimer of Jun and Fos (Rios-Barrera and Riesgo-Escovar, 2013). We present evidence that 20E signaling acts in parallel to the JNK cascade in regulating Jun through activation of EcR, which carries Jun from the cytoplasm to genomic regions containing AP-1 binding sites but no ecdysone-response elements (EcREs) in DC genes. To our knowledge, this the first time that EcR has been shown to directly interact with AP-1 in *Drosophila*, though a genetic interaction has been recently uncovered during the pruning of sensory neuron dendrites (Zhu et al., 2019). Our work demonstrates a mechanism for fine tuning the output from the JNK cascade during DC, and reveals an alternative mode of action for 20E signaling that likely functions beyond DC, as several mammalian steroid hormone receptors can also regulate gene expression in a complex with AP-1 (reviewed in Marino et al., 2006).

## RESULTS

### Dpp signaling to the amnioserosa leads to 20E production, which is required for correct morphogenesis of the tissue during DC

Given that Dpp signaling to the amnioserosa is required for morphogenesis during DC, and that 20E required for DC is produced in the amnioserosa, we tested the hypothesis that Dpp regulated 20E production. An attractive mechanism for the timely production of 20E in the amnioserosa could be through the presence of all but one of the biosynthetic pathway members within the amnioserosa anlage. According to this model, ecdysone production could be activated specifically in the amnioserosa through tissue-specific transcriptional regulation of one or a few of the pathway members. The gene *spook* (*spo*) is the only locus encoding a member of the 20E biosynthetic pathway known to be transcribed in the amnioserosa, although other members of the pathway are expressed in the amnioserosa anlage (Ono et al., 2006). In *tkv^7^* mutant embryos, *spo* expression detected by FISH was largely abolished (Fig. 1A,B). If 20E is required for morphogenesis during DC, then mutants in 20E production should show morphogenetic defects. Indeed, live imaging of embryos mutant for *spo* or *disembodied* (*dib*), another enzyme in the 20E biosynthetic pathway, revealed abnormalities in amnioserosa morphogenesis and a failure to complete DC properly (Fig. 1C-H and Movies S1-S3). In particular, mutants lacking 20E showed uneven contractility of the amnioserosa cells and a failure to complete amnioserosa morphogenesis, suggesting perturbation of cytoskeletal regulation. Thus, candidate genes for regulation by 20E during DC are likely regulators or components of the cytoskeleton expressed in the amnioserosa.

**Fig. 1:**
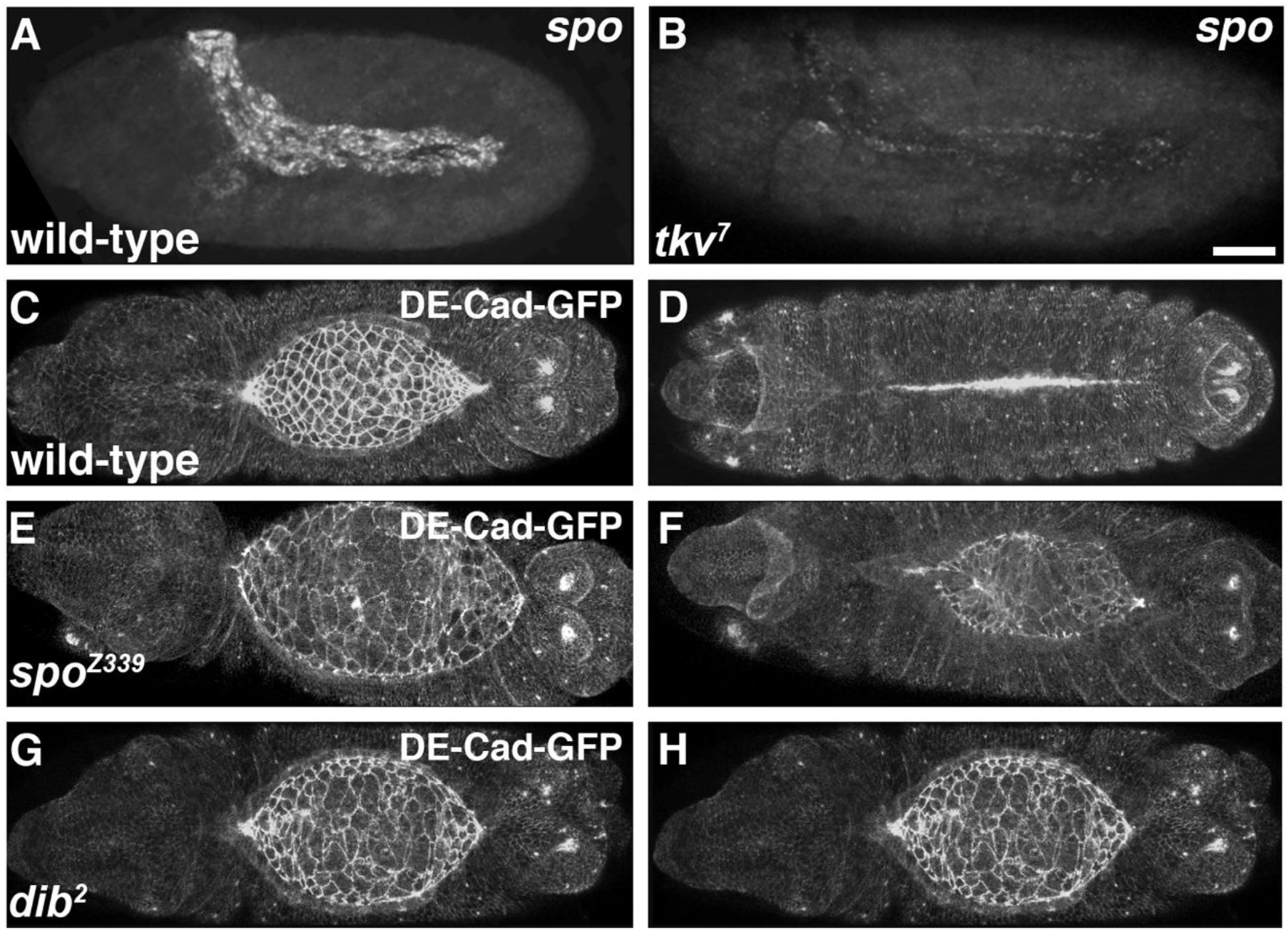
Dpp signaling is required for the expression of *spo*, which, together with another gene involved in 20E biosynthesis, *dib*, is required for correct morphogenesis of the amnioserosa during DC. (A) FISH showing *spo* expression in the amnioserosa during germband retraction in wild-type. (B) In embryos mutant for the Dpp receptor, Tkv, *spo* expression is lost. (C-H) Stills from live imaging of wild-type (C,D), *spo* mutant (E,F), and *dib* mutant (G,H) embryos, showing uniform amnioserosa morphogenesis and closure of the epidermis in wild-type (see Movie S1), but defective amnioserosa morphogenesis and failure of DC in *spo* and *dib* mutant embryos (see Movies S1 and S2). Scale bar represents 50μm (B).

### The timing of expression of four JNK-responsive genes in the amnioserosa is regulated by 20E signaling

Three JNK-responsive genes were previously found to be expressed at high levels in the DME cells and amnioserosa during DC: *jaguar* (*jar*), *jupiter* (*jup*), and *Z band alternatively spliced PDZ-motif protein 52* (*zasp52*) (Ducuing et al., 2015). We determined by FISH that this distribution came from transcriptional regulation and that the duration of expression varied from gene to gene (Fig. S1A-L). *jar* and *zasp52* have been shown to be required for scar-free DC (Ducuing and Vincent, 2016; Millo et al., 2004), while *jup* encodes for a little-studied microtubule-associated protein (Karpova et al., 2006). *zipper* (*zip*) encodes for non-muscle myosin required for cell shape change during DC and is expressed in a similar pattern to these three genes (Fig. S1M-P) (Franke et al., 2005; Young et al., 1993; Zahedi et al., 2008). To test if *zip* was also a JNK-responsive gene, *prd-GAL4* was used to drive embryonic expression of either an activated version of the small Rac1 GTPase, which activates the JNK pathway (Glise and Noselli, 1997; Hou et al., 1997), or a constitutively active form of JNKK encoded by *hemipterous* (*hep*) (Weber et al., 2000). Ectopic expression of Rac1V12 or Hep^CA^ resulted in elevated *zip* transcripts in *prd* stripes in both the epidermis and amnioserosa (Fig. S1Q-S), indicating regulation of *zip* expression by JNK signaling. We confirmed that endogenous JNK signaling was required for this process by impairing the pathway through expression of Bsk^DN^, a dominant negative form of JNK encoded by *basket* (*bsk*) (Weber et al., 2000), which resulted in a loss of *zip* transcripts in *prd* stripes in the DME cells (Fig. S1T).

We next used FISH to examine the expression patterns of the four JNK-responsive genes in embryos mutant for either *spo* or *dib* to determine if loss of 20E also had an effect on their transcription. *jar* and *zasp52* expression normally disappeared from the amnioserosa by the beginning of DC in *spo^1^* and *dib^2^* heterozygous mutant embryos (Fig. 2A,D), which served as controls that displayed similar expression patterns to wild-type (Fig. S1C,K). However, expression of both genes persisted in a subset of amnioserosa cells in *spo^1^* and *dib^2^* homozygous mutant embryos undergoing DC (Fig. 2B,C,E,F). Effects in the DME cells were not readily observable. In contrast to *jar* and *zasp52*, *jup* and *zip* expression in the amnioserosa was shut off earlier in *spo^1^* and *dib^2^* homozygous mutants (Fig. 2H,I,K,L) than in controls (Fig. 2G,J; Fig S1G,M for wild-type). A small decrease in *jup* expression within the DME cells was also observed in the mutants. Quantification of FISH signal can be found in the supplementary material (Fig. S2). Based on these results, we conclude that 20E signaling regulates the timing of the expression of at least four JNK-responsive genes in the amnioserosa during DC.

**Fig. 2:**
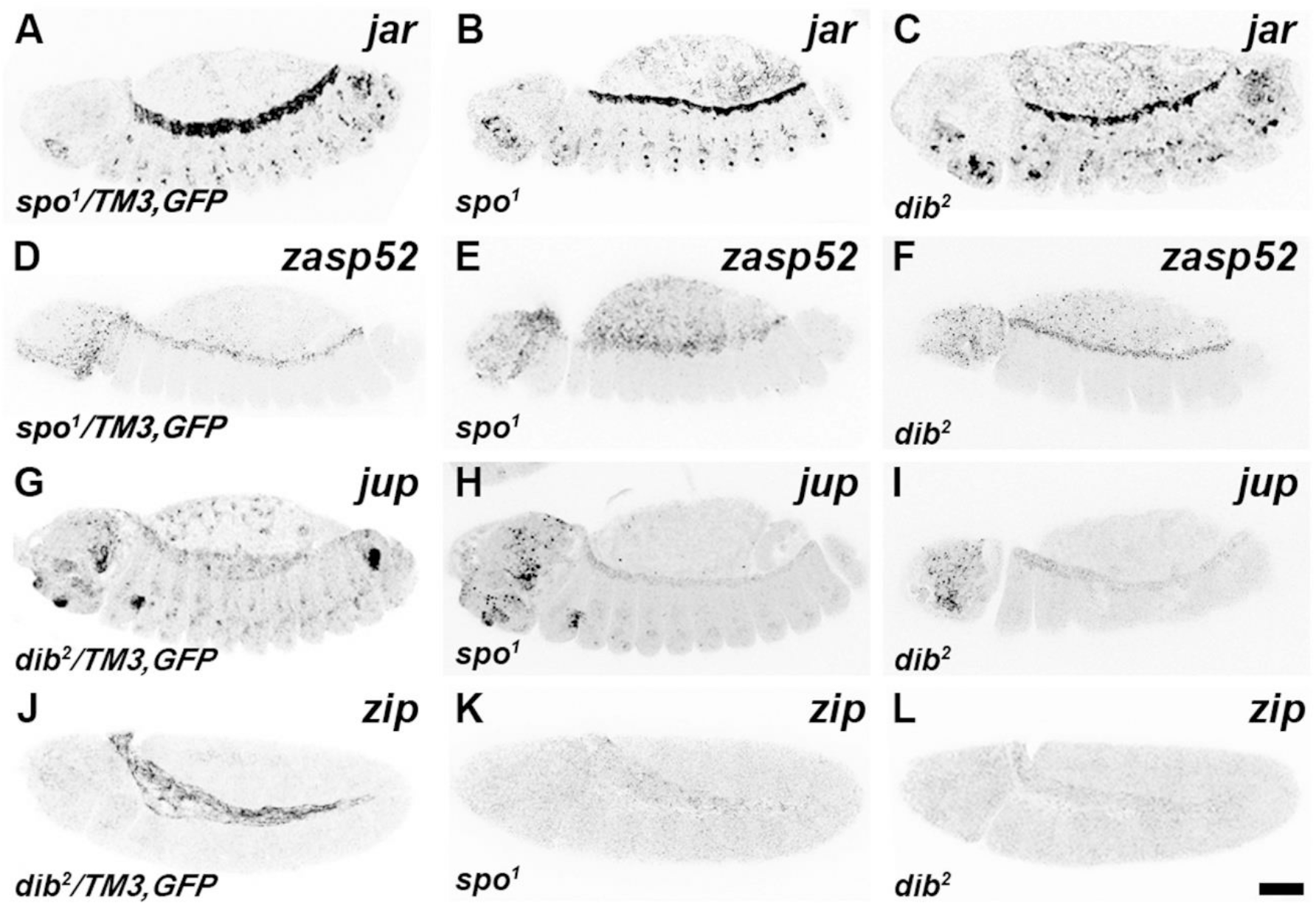
20E signaling regulates the expression of JNK-responsive genes in amnioserosa and DME cells. For clearer views of the changes in gene expression, the images have been inverted. Heterozygous siblings of the homozygous mutant embryos served as controls for each FISH stain, as they were treated under identical conditions within the same tube. (A-C) *jar* expression in the amnioserosa shuts off by the start of DC in the control (A), but persists in a subset of amnioserosa cells in both *spo* and *dib* mutants (B,C) (see Fig. S2A,B for quantifications). Effects in the DME cells were not readily observable (data not quantified). (D-F) Similar results were observed for *zasp52* (see Fig. S2C,D for quantifications). (G-I) Expression of *jup* persists in the amnioserosa during DC in the control (D), but is significantly reduced in both *spo* and *dib* mutants (H,I) (see Fig. S2E,G for quantifications). A slight but significant decrease in expression within the DME cells was also observed in the mutants (see Fig. S2F,H for quantifications). (J-L) *zip* is strongly expressed in the amnioserosa during germband retraction (J), but is lost in both *spo* and *dib* mutants (data not quantified). Scale bar represents 50μm (L).

### EcR forms a complex with the AP-1 component, Jun, in the amnioserosa

20E acts through Ecdysone receptor (EcR), which forms a heterodimer with the nuclear receptor, Ultraspiracle (Usp), and binds to EcREs in target genes (Dobens et al., 1991; Yao et al., 1993). EcR is structurally similar to the vertebrate estrogen receptor, which has been shown to be able to bind to AP-1, the transcription factor acting in the JNK cascade that is commonly composed of heterodimers of Jun and Fos (Marino et al., 2006). Interestingly, JNK signaling is shut off in the amnioserosa prior to DC, with Fos adopting a largely cytoplasmic distribution but with Jun retaining some nuclear localization (Reed et al., 2001). This downregulation of JNK signaling in the amnioserosa is required for DC, and we wondered if there might be a “handing over of control” of gene expression in the amnioserosa from the JNK pathway to ecdysone signaling through an interaction between EcR and Jun. In wild-type, *jar* and *zasp52* lose amnioserosa expression before the completion of germband retraction (Fig. S1B,J), whereas expression of *jup* and *zip* persist longer in the tissue (Fig. S1F,N). We used Bsk^DN^ expression in the amnioserosa to test for a requirement for JNK signaling in maintaining *jup* and *zip* transcription and found that it was not required (Fig. 3A,B). A dominant negative version of EcR, EcR-W650A, which is thought to block endogenous EcR from dimerizing with Usp and thereby repress expression at ecdysone response elements, also failed to block amnioserosa transcription of *jup* and *zip* (Fig. 3C,D), but did block embryonic epidermal transcription of a known ecdysone-responsive gene, *IMP-L1* (Fig. 3E-F) (Cherbas et al., 2003; Natzle et al., 1988; Natzle et al., 1992). These results indicate that expression in the amnioserosa, at least for *jup* and *zip*, is not dependent on JNK or canonical 20E signaling.

**Fig. 3:**
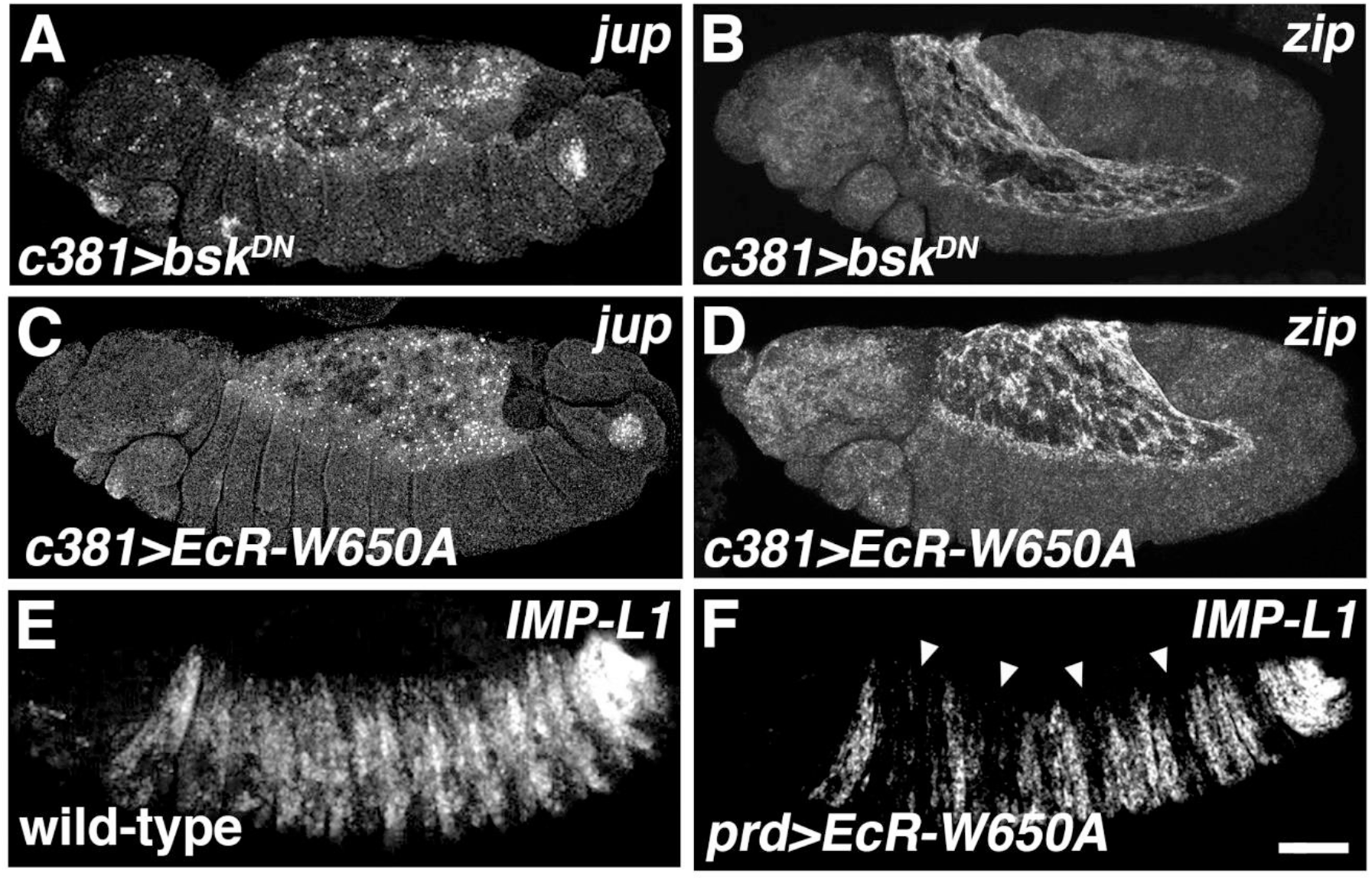
20E-mediated gene expression in the amnioserosa is independent of the JNK and canonical ecdysone pathways. (A,B) Impairment of the JNK pathway in the amnioserosa through Bsk^DN^ expression does not block the transcription of either *jup* (A) or *zip* (B) during germband retraction. (C-F) Impairment of canonical ecdysone signaling in the amnioserosa through the expression of EcR-W650A also does not block *jup* (C) or *zip* (D) transcription, but does block transcription of the known ecdysone-responsive gene, *IMP-L1*, when EcR-W650A is expressed in epidermal *prd* stripes (arrows) (E,F). Scale bar represents 50μm (F).

We wondered if 20E regulates gene expression in the amnioserosa by modulating an interaction between EcR and the AP-1 subunit, Jun, given Jun’s persistent nuclear localization in the tissue. We looked for such an *in vivo* interaction using PLA (Soderberg et al., 2006), and found that Ecr formed a complex with Jun predominately in the amnioserosa from germband retraction to DC (Fig. 4A,B). PLA signal was largely absent in *spo^1^* mutant embryos, indicating a loss of EcR-Jun complexes (Fig. 4C). For these experiments, we used antibodies against EcR and Jun that revealed their presence in amnioserosa nuclei, as well as in the epidermis (Fig. 4D,E). Higher magnification views showed that complexes of EcR and Jun were largely found in amnioserosa nuclei, with much lower levels found throughout the epidermis (Fig. 4E’-G’). Negative controls that were performed with anti-EcR antibody omitted or with anti-Jun replaced by anti-pMad, which detects another transcription factor participating in DC (reviewed in Affolter et al., 2001), showed very low signal background (Fig. S3). It is not surprising that multiple PLA signals are seen in the amnioserosa, as the amnioserosa is the site of high levels of 20E and EcR in the nucleus. We next assessed if the association between Ecr and Jun involved direct physical interaction using reciprocal GST pull-down assays and found that EcR could bind directly to Jun, *in vitro* (Fig. 4H,I). Interestingly, these assays also showed that EcR could bind to Fos (Kayak, Kay, in *Drosophila*) and Jun could bind to Usp, though further work is required to confirm the *in vivo* relevance of these interactions.

**Fig. 4:**
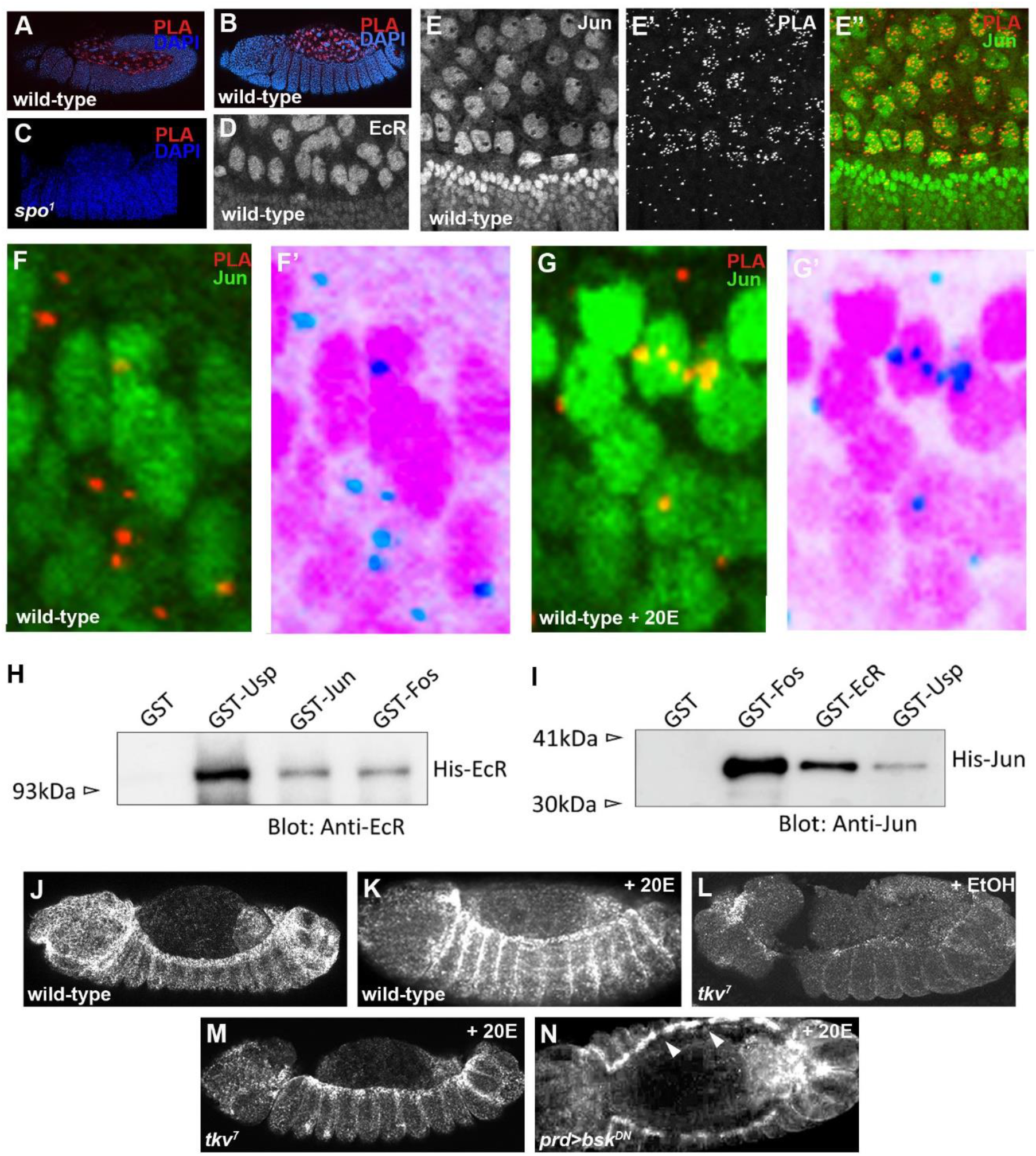
Evidence of interactions between 20E signaling and the JNK pathway. (A,B) Wild-type embryos subjected to PLA between EcR/Jun (red) and stained with DAPI (blue) predominately show clusters of PLA complexes in amnioserosa nuclei during germband retraction (A) and DC (B). (C) PLA signals are not observed in *spo* mutant embryos. (D) Close-up view of a wild-type embryo stained with anti-EcR antibody shows highest levels of EcR in amnioserosa nuclei during DC (this antibody was used in the PLA experiments). (E-E’’) Wild-type embryo subjected to PLA between EcR/Jun (red in E’,E’’) and stained with anti-Jun antibody (E, green in E’’). Highest levels of Jun are found in the DME cells, but Jun is also present in amnioserosa nuclei during DC. (F-G’) Similarly stained embryos as in E-E’’. High magnification views of epidermal cells show that PLA complexes are largely cytoplasmic in wild-type embryos (F,F’), where endogenous 20E levels are low, but translocate into the nucleus upon 20E-treatment (G,G’). F’ and G’ are inverted images. (H,I) Immunoblot analysis of pull-down assays between EcR and Jun. EcR immunoblots show that GST-Jun and GST-Fos (Kay) are both able to pull-down His-EcR (H). No binding was observed in the negative control, which involved GST alone. GST-Usp served as a positive control since Usp is known to dimerize with EcR. Jun immunoblots show that GST-EcR and GST-Usp were both able to pull-down His-Jun in reciprocal assays (I). No binding was observed with GST alone. GST-Fos (Kay), a component of the AP-1 transcription factor, served as a positive control. (J,K) Wild-type embryos treated with 20E show ectopic *zip* transcription in the epidermis (K) when compared to untreated embryos (J). (L,M) *tkv* mutant embryos show reduced *zip* transcript levels (L), but upon 20E-treatment, *zip* transcription is restored in the DME cells (M). (N) In contrast, 20E-treatment does not restore *zip* transcription in DME cells expressing Bsk^DN^ (arrowheads).

In the embryonic epidermis, where 20E levels are lower, complexes of EcR and Jun were also observed but were consistently outside the nucleus, with 72.5% of 131 PLA signals counted in a wild-type embryo being cytoplasmic (Fig. 4F,F’). Soaking embryos in 20E caused EcR-Jun complexes in the epidermis to translocate into the nucleus with only about a third of PLA signals remaining in the cytoplasm (Fig. 4G,G’), and this was accompanied by elevated expression of *zip* transcripts in the epidermis (Fig. 4J,K). Collectively, these results suggest that high levels of 20E promote the movement of EcR-Jun complexes into the nucleus where they can modulate gene expression.

### 20E signaling requires the JNK pathway to drive ectopic *zip* expression in the epidermis

Having determined that exogenous 20E can elevate *zip* expression in the embryonic epidermis, we explored the requirements for such regulation. We first assessed the ability of exogenous 20E to restore *zip* transcription in the DME cells of *tkv^7^* mutant embryos, in which endogenous 20E is absent, and found that it could (Fig. 4L,M). As seen above, knockdown of the JNK pathway in the amnioserosa through expression of Bsk^DN^ did not prevent 20E-dependent gene expression in that tissue. In contrast, exogenous 20E was incapable of restoring *zip* transcription in DME cells with Bsk^DN^ expression (Fig. 4N), indicating a requirement for JNK pathway activation in triggering 20E-induced ectopic expression of *zip* in the epidermis.

### Discovery of putative EcR-AP-1 enhancers in the introns of DC genes

We have shown that 20E is required for the expression of *zip* in the amnioserosa - is there any evidence of EcR directly binding to the *zip* locus? Gauhar and colleagues mapped 502 genomic binding regions for EcR-Usp in *Drosophila* Kc167 cells treated with 20E, one of which resides in intronic sequences of *zip* (Fig. 5A) (Gauhar et al., 2009). This region lacks a consensus EcRE but does contain five copies of the AP-1 binding motif consensus, TGANTCA, suggesting EcR binding to the *zip* locus through its association with Jun. We wondered if this region constituted an enhancer modulating gene expression in the amnioserosa by EcR and Jun and screened through the KcI67 EcR-Usp binding regions for those containing at least four consensus AP-1 binding motifs but no EcRE consensus site. We identified 49 additional genomic regions fitting these criteria, 20% of which were strikingly in or near known DC genes including those encoding the transcription factors Cabut (Cbt) and U-shaped (Ush), the non-receptor tyrosine kinase Activated Cdc42 kinase (Ack), the receptor tyrosine kinase Insulin-like receptor (InR), the G protein-coupled receptor kinase Gprk2, the Arf-GEF Steppke (Step), the cytoskeletal regulators Short stop (Shot), Enabled (Ena) and Rho1, and EcR itself; this latter finding suggesting a feedback loop (see Table 1 for more information) (Fernandez et al., 1995; Grevengoed et al., 2001; Harden et al., 1999; Lada et al., 2012; Munoz-Descalzo et al., 2005; Schneider and Spradling, 1997; Sem et al., 2002; Takacs et al., 2017; West et al., 2017). In an effort to look for further evidence of joint regulation of EcR and Jun in such genes, we used ChIP-seq data generated by Kevin White’s lab as part of the ENCODE consortium (Davis et al., 2018; ENCODE Project Consortium, 2012). These data include genome-wide binding regions for GFP-tagged versions of EcR, Usp, Jun and Fos, immunoprecipitated with anti-GFP antibodies from white prepupae, 0-12 hour old embryos, wandering third instar larvae, and 0-24 hour old embryos, respectively. Putative binding regions for these proteins were scattered throughout *zip* introns, but not in open reading frames (Fig. 5A).

**Fig. 5:**
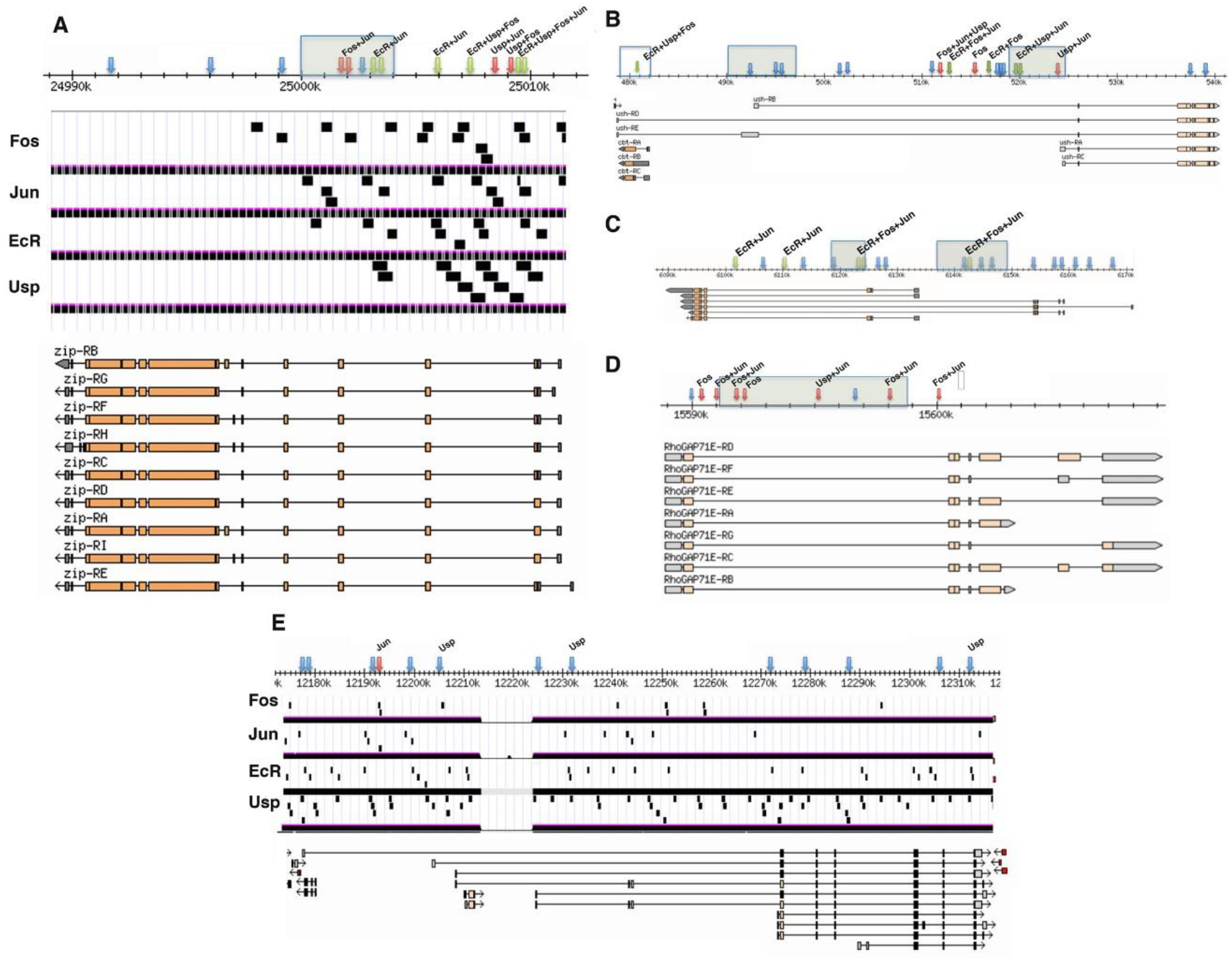
Putative EcR-AP-1 enhancers are located in large introns of genes expressed in dorsal tissues during germband retraction and DC. Diagrams are modified from GBrowse and UCSC Genome Browser. Arrows mark consensus AP-1 binding sites (TGANTCA). Blue arrows are sites that do not overlap with ChIP-seq peaks for EcR, Jun or Fos; red arrows are sites that overlap with ChIP-seq peaks for Jun and/or Fos; green arrows are sites that overlap with ChIP-seq peaks for EcR. Labels above arrows indicate ChIP-seq peaks that contain consensus AP-1 binding sites, whereas shaded boxes are EcR binding regions identified by Gauhar and colleagues (Gauhar et al., 2009). Sample distributions of ChIP-seq peaks are denoted in panels A and E as black rectangles. (A) *zip* locus showing no binding of the four transcription factor proteins to exons. (B) *cbt* and *ush* genomic region. The unshaded box on the far left denotes sequences controlling *cbt* expression. (C) *EcR* genomic region. (D) *RhoGAP71E* genomic region. (E) Control large intron gene, *bru*, showing distribution of consensus AP-1 binding sites in a gene not known to be regulated by JNK or 20E signaling. There is only about one AP-1 binding site every 10kb.

To explore further the regulation of gene expression by EcR acting at AP-1 binding sites, we selected five genes from the screen to determine if 20E regulates their expression with FISH. The five genes were the known DC participants *cbt* and *ush*, plus *EcR*, *RhoGAP71E* and *Mes2*. All of these genes are expressed in the amnioserosa (Fig. S4), and enriched in other dorsal tissues including the yolk sac and hindgut (*cbt*, Fig. S4A-C), the dorsal epidermis (*ush*, Fig. S4D-F), and the dorsal vessel (*RhoGAP71E*, Fig. S4J-L; *Mes2*, Fig. S4M-O) (Belacortu et al., 2011; Kozlova and Thummel, 2000; Lada et al., 2012).

*cbt* is located in an intron of *ush*, but transcribed in the opposite direction. Based on previous immunostains, Cbt is expressed in yolk sac nuclei, the amnioserosa, as well as in other more ventral tissues during DC (Belacortu et al., 2011). Expression in the yolk sac and amnioserosa appeared unperturbed in *spo^1^* and *dib^2^* mutant embryos, but relative to these two tissues, *cbt* transcript levels were elevated in the epidermis (Fig. 6A-D; quantifications in Fig. S5A-D), indicating inhibition of *cbt* expression by 20E signaling. A previous study used reporter gene expression studies to identify a block of sequences that promoted gene expression in many of the tissues Cbt is found and likely constitutes the major control region for *cbt* (Belacortu et al., 2011). This region has a single AP-1 binding motif, which EcR and Fos have been shown to bind in the vicinity of (Fig. 5B). Starting about 8kb upstream of the *cbt* regulatory region is a stretch of about 35kb of intronic sequences with multiple consensus AP-1 binding motifs that putatively recruit various combinations of Ecr, Usp, Jun and Fos, and are likely control sequences for *ush* expression. We found that the expression of *ush* in the peripheral amnioserosa cells and dorsal epidermis were reduced in *spo^1^* and *dib^2^* mutant embryos (Fig. 6E-H; quantifications in Fig. S5E-H), indicating promotion of *ush* expression by 20E signaling.

**Fig. 6:**
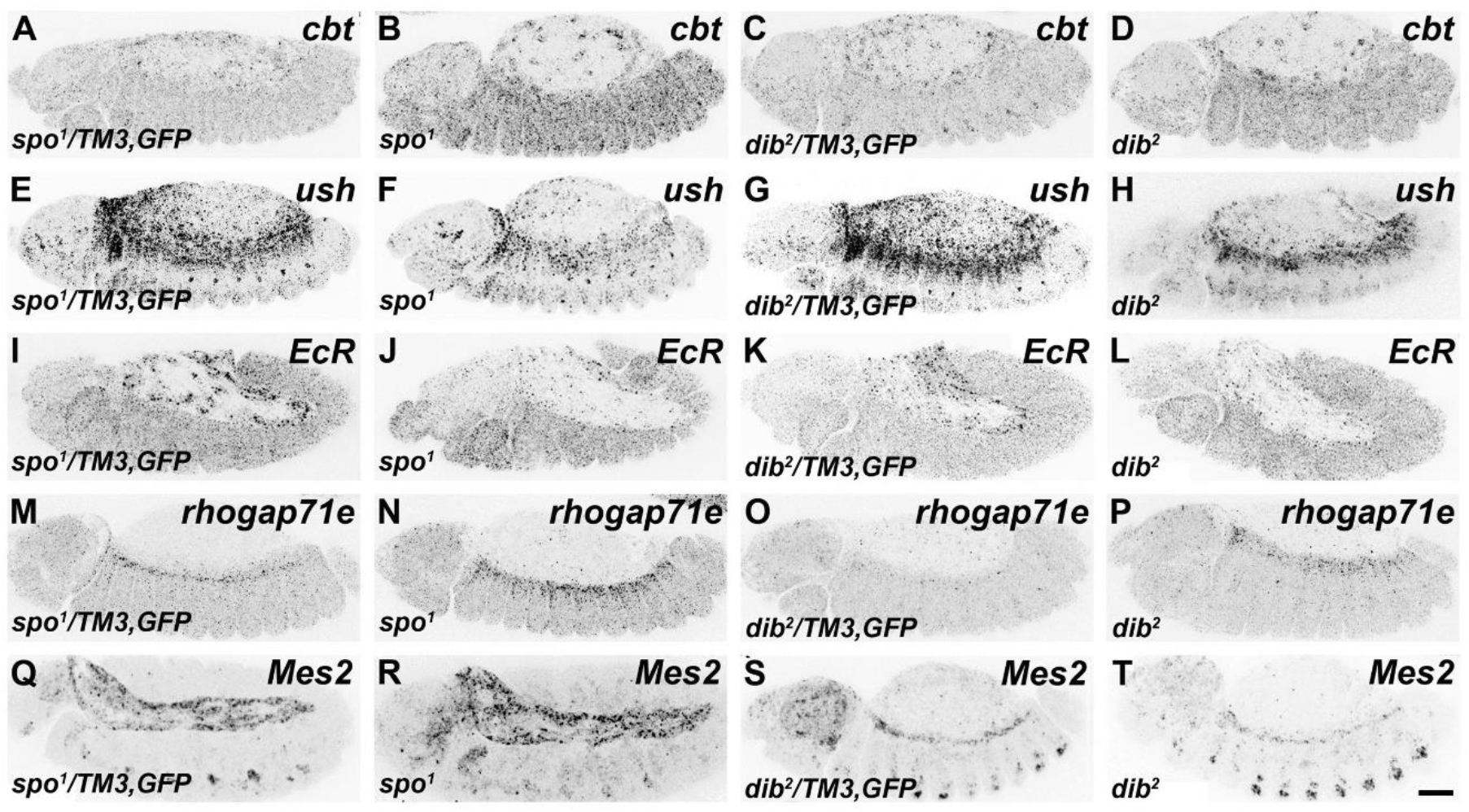
Loss of 20E signaling affects the expression of genes bearing putative EcR-AP-1 enhancers. For clearer views of the changes in gene expression, the images have been inverted. Heterozygous siblings of the homozygous mutant embryos served as controls for each FISH stain, as they were treated under identical conditions within the same tube. (A-D) Relative to the controls (A,C), *spo* and *dib* mutant embryos both show increased *cbt* expression in the epidermis but no change in the amnioserosa (B,D) (see Fig. S5A-D for quantifications). (E-H) Control embryos have high levels of *ush* expression in both the peripheral amnioserosa cells and dorsal epidermis (E,G), but expression is reduced in both tissues of embryos mutant for either *spo* or *dib* (F,H) (see Fig. S5E-H for quantifications). (I-L) In contrast to the controls (I,K), expression of *EcR* in the amnioserosa during germband retraction is lost with disruption of 20E signaling (J,L) (see Fig. S5I,J for quantifications). (M-P) *RhoGAP71E* expression is restricted to the dorsal vessel in control embryos (M,O), but is ectopically expressed in the dorsal epidermis in both *spo* and *dib* mutant embryos (N,P) (see Fig. S5K,L for quantifications). (Q-T) No change in the expression of *Mes2* is observed between control (Q,S) and mutant (R,T) embryos (see Fig. S5M-P for quantifications). Scale bar represents 50μm (T).

Consensus AP-1 binding motifs were found in four binding regions for EcR in the *EcR* locus, including two in the EcR-bound area found in 20E-treated Kc167 cells (Fig. 5C). All these AP-1 motifs were also found in binding regions for Jun and/or Fos, supporting the idea that EcR is guided to binding sites in a complex with AP-1. *EcR* expression comes on strongly in the amnioserosa during germband retraction in wild-type but not in *spo^1^* nor *dib^2^* mutant embryos (Fig. 6I-L; quantifications in Fig. S5I,J), suggesting that EcR operates in a positive feedback loop for 20E-mediated gene expression in the amnioserosa. *RhoGAP71E* expression in wild-type is typically restricted to the dorsal vessel during DC, but expression was ectopically induced in the dorsal epidermis of *spo^1^* and *dib^2^* mutants (Fig. 6M-P; quantifications in Fig. S5K,L). Interestingly, although an EcR-bound region within the *RhoGAP7E* locus of 20E-induced Kc167 cells was identified, the ENCODE consortium data showed no binding of EcR to this region, but one instance of Usp binding (Fig. 5D). Finally, despite *Mes2* being isolated as a putative EcR-binding gene with AP-1 sites, loss of 20E had no discernible effects on *Mes2* expression in the amnioserosa or the dorsal vessel (Fig. 6Q-T; quantifications in Fig. S5M-P).

We noticed that the consensus AP-1 binding motifs apparently recruiting EcR and the AP-1 components to DNA tended to occur in large introns. Indeed, the candidate genes from our search (listed in Table 1) were on average twice the size of the average *Drosophila* gene (*i.e.* 22kb compared to 11kb). This suggested that long stretches of unconstrained DNA could be the raw material for the emergence of these enhancers and we examined the distributions of consensus AP-1 binding motifs and ChIP data for several large genes including *brn-1* (Fig. 5E). Such genes had many ChIP-seq peaks scattered throughout their introns that showed little overlap with the AP-1 binding motifs, suggesting that many of the ChIP-seq peaks represent spurious interactions and/or binding to non-consensus sequences (Spivakov, 2014).

## DISCUSSION

With amnioserosa morphogenesis being a critical component of DC, it is important that the timing and degree of cell constriction is carefully modulated and is synchronized with morphogenesis of the epidermis. We have provided evidence here that Dpp signaling from the epidermis informs the amnioserosa that epidermal morphogenesis is commencing by turning on *spo*. This leads to 20E production in the amnioserosa, which can then regulate gene expression in the amnioserosa and nearby tissues such as the dorsal epidermis by promoting complexes of EcR with the AP-1 component, Jun, and carrying it into the nucleus. The most commonly regulated tissue is the amnioserosa, with six of nine genes examined in this study showing modulation by 20E. This is not surprising as the amnioserosa has the highest levels of EcR, 20E, and EcR-Jun complexes during germband retraction and DC.

We see three patterns of wild-type gene expression in the amnioserosa, which may reflect differing contributions from JNK and 20E signaling. The first pattern, seen with *RhoGAP71E* and *Mes2*, which is likely driven predominately by JNK but only weakly, is modest gene expression prior to germband retraction that quickly disappears as germband retraction begins, presumably as the JNK pathway is shut down in the amnioserosa. An effect of 20E signaling on this expression is not apparent. The second pattern, seen with *jar* and *zasp52*, is intense expression before the start of germband retraction that is driven by JNK, but subsequently shuts down during germband retraction. This requires EcR-mediated repression, such that loss of 20E signaling results in persistence of gene expression in the amnioserosa into DC. The third pattern, seen with *jup*, *zip* and *EcR*, is persistence of gene expression throughout germband retraction that requires 20E signaling. In this situation, there may be a “handing off” of gene expression whereby EcR takes over control of AP-1 as the JNK pathway shuts down in the amnioserosa. EcR does not normally promote gene expression in the epidermis, which is JNK-dependent, presumably because 20E levels are too low. Indeed, when 20E levels are artificially elevated, *zip* is ectopically expressed. We have also provided evidence of gene repression by 20E signaling in the epidermis, as seen with *cbt* and *RhoGAP71E*.

Our data indicate that EcR can both positively and negatively regulate expression of AP-1-dependent genes. Similarly, estrogen activates some genes through AP-1 and represses others (Bjornstrom and Sjoberg, 2005). Future work will be aimed at determining the composition of EcR complexes and how they activate or repress gene expression. We have not established if the EcR partner Usp or the Fos subunit of AP-1 are involved, although ChIP-seq data and our GST pulldowns suggests they could part of the complex binding to these enhancers, which we will refer to as EcR-AP-1 enhancers.

We have demonstrated that 20E signaling, acting through EcR-AP-1 enhancers, allows for more refined modulation of gene expression during DC than the JNK pathway on its own. Presumably these enhancers are acting elsewhere during development when and where 20E signaling and the JNK cascade overlap, and this is supported by ChIP-seq data suggesting spatial and temporal variation in enhancer binding. A good candidate tissue is the salivary gland, where AP-1 is required for 20E-triggered cell death (Lehmann et al., 2002).

An important question raised by our data is do they tell us about the origin of steroid hormone-AP-1 interactions? The large introns of the genes containing the EcR-AP-1 enhancers may have provided an ideal setting for the emergence of these regulatory sequences by allowing transcription factors to experiment with their DNA binding, which could be followed by the evolution of protein-protein interactions between transcription factors fortuitously finding themselves as neighbors on DNA. This could be a mechanism for convergent evolution of steroid hormone receptor interactions. In the absence of molecular comparisons between *Drosophila* and vertebrate steroid hormone receptor-AP-1 complexes, it is uncertain if our results support an ancient origin of interactions between these transcription factor families.

## MATERIALS AND METHODS

### Fly stocks

Flies were maintained at 25°C under standard conditions (Ashburner and Roote, 2007). *w^1118^* was used as a wild-type control strain unless otherwise stated. *spo^Z339^* was a kind gift from M. O’Connor (Ono et al., 2006), and *ubi-DE-cadherin-GFP* was generously provided by H. Oda (Oda and Tsukita, 2001). All other stocks were obtained from the Bloomington *Drosophila* Stock Center.

### Live imaging of embryos

Embryos were prepared for live imaging using the hanging drop protocol (Reed et al., 2009), and imaged with a Nikon Eclipse 90i microscope with a Nikon D-Eclipse C1 scan head. Images were saved as animated projections using Nikon EZ-C1 software and further processed with ImageJ (NIH). A *ubi-DE-cadherin-GFP* transgene was expressed in all embryos to visualize morphology (Oda and Tsukita, 2001).

### 20E-treatment of embryos

Embryonic treatment with exogenous 20E was performed as previously described (Kozlova and Thummel, 2003). Embryos were collected for six hours (Rothwell and Sullivan, 2007a), then cultured for another four hours in MBIM, supplemented with 5×10^−6^M 20E (H5142, Sigma-Aldrich) dissolved in ethanol, prior to fixation (Rothwell and Sullivan, 2007b). Control embryos, done in parallel, were subjected to the same treatment but replacing 20E in ethanol with ethanol alone.

### Fluorescent *in situ* hybridization (FISH)

Detection of transcripts *in situ* by FISH was performed as described (Lecuyer et al., 2007). cDNA templates used to make full-length antisense probes were obtained from the *Drosophila* Genomics Resource Center. Fluorescently-stained embryos were examined on a Nikon A1R laser scanning confocal microscope with NIS-Elements software, and the images were processed with Adobe Photoshop. Mutant stocks were rebalanced over GFP-tagged balancers allowing for homozygotes to be selected based on the absence of GFP signal. Heterozygous siblings, which were treated under identical conditions within the same tube, served as controls. For transgenic analysis, homozygous *UAS*-transgene-bearing males were crossed to homozygous *Gal4*-bearing virgin females ensuring that all progeny carried one copy of each. In cases where either the *Gal4* or *UAS*-transgenic stock was homozygous lethal, the stock was also re-balanced over a GFP-containing balancer. In subsequent crosses, GFP-negative embryos carried both the *Gal4* and *UAS*-transgene, whereas GFP-positive embryos lacked either the *Gal4* or *UAS*-transgene and, therefore, had no transgenic expression.

### Quantification of FISH signal

Quantification of FISH signal from stacked confocal images is described in the supplementary material.

### Proximity ligation assay (PLA)

PLA was performed as previously described but with modifications (Thymiakou and Episkopou, 2011). Fixed embryos (Rothwell and Sullivan, 2007a; Rothwell and Sullivan, 2007b) were blocked for one hour with 1% BSA (in PBT: 3mM NaH^2^PO^4^·H_2_O, 7mM Na_2_PO_4_, 1.3M NaCl, 0.1% Triton X-100, pH 7.0). Next, the embryos were incubated with 1:5 mouse anti-EcR (DDA2.7, Developmental Studies Hybridoma Bank) (Talbot et al., 1993) and 1:25 rabbit anti-Jun (sc-25763, Santa Cruz Biotechnology) primary antibodies in 1% BSA overnight at 4°C. After three PBT washes for ten minutes each, the embryos were incubated with 1:5 dilutions of anti-rabbit PLUS (DUO92002, Sigma-Aldrich) and anti-mouse MINUS (DUO92004, Sigma-Aldrich) PLA probes in 1% BSA for two hours at 37°C. The embryos were subsequently washed twice with Wash A for five minutes each, then incubated in Ligation reagent (DUO92008, Sigma-Aldrich) for one hour at 37°C. Following two washes with Wash A for two minutes each, the embryos were incubated in Amplification reagent (DUO92008, Sigma-Aldrich) for two hours at 37°C. After two Wash A washes for two minutes each, the embryos were incubated with 1:200 FITC-conjugated anti-mouse or anti-rabbit secondary antibody (Jackson ImmunoResearch) in 1% BSA for one hour. Finally, the embryos were washed twice with Wash B for ten minutes each, followed by a single wash with 0.01x Wash B for one minute, then stored in Duolink In Situ Mounting Medium with DAPI (DUO82040, Sigma-Aldrich) at −20°C until ready for confocal imaging.

### GST pull-down assays

Preparation of tagged proteins was performed as previously described (Rebay and Fehon, 2009). The following cDNA clones, obtained from the *Drosophila* Genomics Resource Center, were used: *EcR* (RE06878), *jun* (LD25202), *usp* (LD09973), and *kay* (LP01201). Full-length coding regions were amplified and inserted in frame into pET-28a(+) (69864-3, MilliporeSigma) and/or pGEX-4T-1 (28-9545-49, GE Healthcare) to create N-terminal, His- and GST-tagged constructs, respectively. The constructs were transformed into BL21(DE3) competent cells for expression (C2527, New England Biolabs).

Pull-downs were standardized by adding an equivalent amount of bait protein (*i.e.* the GST-tagged protein from the bacterial soluble protein fraction) to an equal volume of prey protein (*i.e.* the His-tagged protein from the bacterial soluble protein fraction). The volume was then topped up to 500μL with Buffer A (20mM Tris, 1mM MgCl2, 150mM NaCl, 0.1% NP-40, 10% Glycerol, 1x protease inhibitor, pH 8.0), and the mix was incubated for 1.5 hours at 4°C. In the meantime, 25μL of Glutathione Sepharose 4B (17-0756-01, GE Healthcare) was blocked with 1% BSA (in Buffer A) for one hour at 4°C. The mix was then added to the blocked beads and incubated for another 1.5 hours at 4°C. Following three washes with Buffer A, bound proteins were denatured and fractionated by SDS-PAGE. The presence of His-tagged, prey proteins was determined by immunoblotting with the use of the following primary antibodies: 1:150 mouse anti-EcR (DDA2.7, DSHB) (Talbot et al., 1993) and 1:1000 rabbit anti-Jun (sc-25763, SCBT). Both antibodies were diluted in 1% milk (in TBST: 1.5M Tris, 0.5M NaCl, 0.1% Tween 20, pH 7.5). Peroxidase-conjugated secondary antibodies (Vector Laboratories) were used at a 1:2000 dilution in 1% milk, and signal was detected with BM Chemiluminescence Blotting Substrate (11500694001, Roche).

## ACKNOWLEDGMENTS

We thank M. O’Connor and H. Oda for kindly gifting fly lines. Stocks obtained from the Bloomington *Drosophila* Stock Center (NIH P40OD018537) were used in this study. cDNA clones were obtained from the *Drosophila* Genomics Resource Center (NIH 2P40OD010949). DDA2.7 (EcR common) antibody, deposited by C. Thummel and D. Hogness, was obtained from the Developmental Studies Hybridoma Bank, created by the NICHD of the NIH and maintained at the University of Iowa.

